# *Chst9* Marks a Spatially and Transcriptionally Unique Population of *Oprm1*-Expressing Neurons in the Nucleus Accumbens

**DOI:** 10.1101/2023.10.16.562623

**Authors:** Emma Andraka, Robert A. Phillips, Kasey L. Brida, Jeremy J. Day

## Abstract

Opioids produce addictive, analgesic, and euphoric effects via actions at mu opioid receptors (μORs). The μOR is encoded by the *Oprm1* gene and is expressed in multiple brain regions that regulate reward and motivation, such as the nucleus accumbens (NAc). *Oprm1* expression in NAc medium spiny neurons (MSNs) mediates opioid place preference, seeking, and consumption. However, recent single nucleus RNA sequencing (snRNA-seq) studies in rodent, primate, and human NAc have revealed that multiple subpopulations of NAc neurons express *Oprm1* mRNA, making it unclear which populations mediate diverse behaviors resulting from μOR activation. Using published snRNA-seq datasets from the rat NAc, we identified a novel population of MSNs that express the highest levels of *Oprm1* of any NAc cell type. Here, we show that this population is selectively marked by expression of *Chst9*, a gene encoding a carbohydrate sulfotransferase. To validate this observation and characterize spatial localization of this population in the rat NAc, we performed multiplexed RNAscope fluorescence *in situ* hybridization studies to detect expression of *Oprm1* and *Chst9* mRNA along with well-validated markers of MSNs. Notably, *Chst9*+ neurons exhibited more abundant expression of *Oprm1* as compared to other cell types, and formed discrete cellular clusters along the medial and ventral borders of the NAc shell subregion. Moreover, *CHST9* mRNA was also found to mark specific MSN populations in published human and primate snRNA-seq studies, indicating that this unique population may be conserved across species. Together, these results identify a spatially and transcriptionally distinct NAc neuron population characterized by the expression of *Chst9*. The abundant expression of *Oprm1* in this population and the conservation of these cells across species suggests that they may play a key functional role in opioid response and identify this subpopulation as a target for further investigation.

## INTRODUCTION

Opioids induce rewarding and reinforcing properties by acting at endogenous opioid receptors in the central nervous system, and repeated experience with opioids can lead to physical dependence, substance use disorder, and overdose-related deaths. Endogenous and synthetic opioids activate a conserved family of Gα_i_-coupled GPCRs that include the mu, delta, and kappa opioid receptors, which are encoded by *Oprm1, Oprd1*, and *Oprk1*, respectively. Commonly used opioid drugs such as morphine and fentanyl produce reward via actions primarily at the mu opioid receptor (μOR). For example, morphine conditioned place preference and dependence is lost in transgenic mice with constitutive deletion of the *Oprm1* gene^1^, demonstrating that this receptor is required for opioid reward. Similarly, the μOR plays critical roles in motivation for natural rewards and social behaviors. *Oprm1*^-/-^ mice do not increase palatable food consumption after a period of deprivation^2^, and both *Oprm1*^-/-^ and *Oprm1*^+/-^ mice have impaired social interactions^3^. However, developing a complete understanding of μOR contributions to behavior is difficult due to the broad expression of *Oprm1* in many distinct brain nuclei and cell types^4,5^, each of which may contribute to distinct opioid actions.

A major target of investigation for the reward-related functions of opioids is the nucleus accumbens (NAc), a striatal subregion that integrates information about motivational states, hedonic stimuli, reward-paired cues, and contextual information^6–8^. The NAc is divided into two subregions, the core and shell. The core envelops the anterior commissure and is architecturally and functionally similar to the dorsal striatum. The NAc shell represents the most medial and ventral aspect of the NAc. While the NAc core facilitates learning and reward-related associations, the NAc shell is primarily implicated in the rewarding and reinforcing qualities of relevant stimuli^6,7,9^. Ample evidence has also demonstrated that the NAc serves as a hub for the effects of opioids on motivated behavior^5,6^. Selective restoration of *Oprm1* expression in a subset of striatal neurons in *Oprm1*^-/-^ mice rescues morphine reward, without altering the analgesic properties or withdrawal effects of morphine^10^. Likewise, direct infusions of μOR agonists into the NAc increase food consumption^8,11^, sucrose drinking^12^, and intake of high-fat foods^13,14^, whereas intra-Nac μOR antagonists decrease sucrose intake and feeding behavior^15^. Additionally, opioid receptor antagonists delivered into the NAc block opioid reinforcement and social reward^16,17^, and direct infusion of μOR receptor agonists into the NAc is reinforcing in self-administration tasks^16,18,19^.

The principal cells of the NAc are GABAergic medium spiny neurons (MSNs) that directly target the output nuclei of the basal ganglia (ventral tegmental area and substantia nigra) or indirectly target these nuclei through projections to the ventral pallidum^20–22^. MSNs are further separated by expression of the *Drd1* and *Drd2* dopamine receptors, which form distinct MSN populations termed D1-MSNs and D2-MSNs^23,24^. While this division is critical for understanding striatal physiology^25^, circuitry^26^, and molecular responses to drugs of abuse^27–29^, both cell populations express *Oprm1*^30^ and contribute to opioid-induced behaviors^31^. Further, recent single nucleus RNA sequencing (snRNA-seq) studies have revealed unexpected heterogeneity in MSNs within the NAc and broader striatum, with multiple subtypes of *Drd1* and *Drd2* expressing MSNs present in the mouse^32^, rat^33–35^, monkey^36,37^, and human NAc^37,38^. Strikingly, NAc MSN subtypes exhibit highly variable expression of *Oprm1*^34^, highlighting the need for further investigation into the roles of distinct MSN subclasses in opioid action. Here, we leveraged available snRNA-seq datasets to characterize a subpopulation of NAc MSNs that exhibit the highest level of *Oprm1* expression across all defined NAc cell types. Using data-driven comparisons, we identified a marker gene (*Chst9*, which encodes a carbohydrate sulfotransferase) that defines this neuronal population. Next, we employed single molecule RNA fluorescence in situ hybridization to validate this marker and map the spatial location of this transcriptionally defined cell type in the rat NAc. Finally, comparison of transcriptional profiles of MSN subtypes from the rat, monkey, and human NAc revealed that this cell population is conserved across species. Together, these findings highlight a unique population of NAc neurons with abundant *Oprm1* expression and identify a specific molecular marker that may be useful in future targeting of this cell type.

## RESULTS

### Computational identification of a novel marker gene Grm8-MSNs

To begin examining subpopulations of *Oprm1*+ cells in the NAc, we first consulted a previously published snRNA-seq dataset from the rat NAc^34^. This study identified 16 transcriptionally distinct cell populations, including principal MSNs that express *Ppp1r1b* (the gene encoding dopamine- and cyclic AMP-related phosphoprotein-32, or DARPP-32; **Fig. 1a-b**) among other canonical MSN markers (**Fig. 4d**). MSN populations in the rat NAc included previously characterized D1-MSNs and D2-MSNs, as well as a novel *Ppp1r1b*+ population termed Grm8-MSNs due to enrichment of the gene *Grm8*, the gene encoding the metabotropic glutamate receptor 8 (**Fig. 1c**). Interestingly, *Oprm1* mRNA was highly expressed within a large proportion of Grm8-MSNs (**Fig. 1d**). However, we also observed less abundant *Grm8* and *Oprm1* expression in a smaller fraction of D1 and D2-MSN subpopulations (**Fig. 1c-d**), meaning this subpopulation cannot be distinguished by the colocalization of these genes alone.

**Figure 1.**
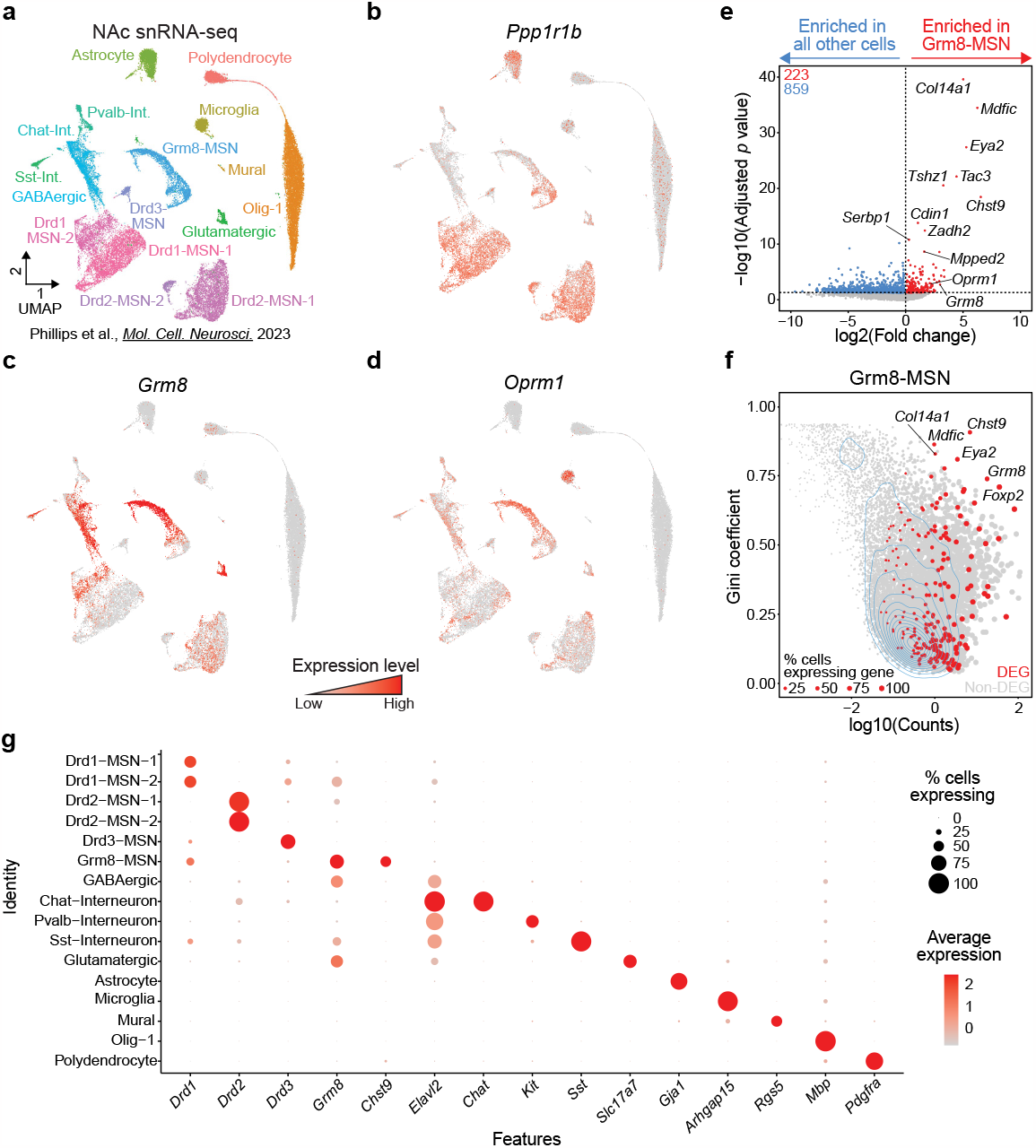
Identification of Chst9 as a marker gene for Grm8-MSNs. **a**, UMAP of previously published snRNA-seq data from NAc of male and female Sprague-Dawley rats identifies 16 transcriptionally distinct cell populations. **b-d**, Feature plots showing distribution of genes expressed in Grm8-MSNs, including *Ppp1r1b* (a pan-MSN marker), *Grm8*, and *Oprm1*. **e**, Volcano plot for identification of differentially expressed genes enriched in Grm8-MSNs. **f**, Gini coefficient analysis identifies *Chst9* as a marker gene with high and selective expression in Grm8-MSNs. **g**, Dot plot for identified marker genes for identified cell populations.

To further investigate the unique gene expression signatures of these Grm8-MSNs, we conducted a differential expression analysis using pseudobulked gene expression matrices across 8 biological replicates. Comparing the gene expression signature of the Grm8-MSN cluster to all other cell clusters identified 223 genes enriched in the Grm8-MSN cluster and 859 genes significantly depleted in this population. Among enriched genes, *Chst9* (encoding carbohydrate sulfotransferase 9) was the most highly enriched differentially expressed gene in the Grm8-MSN population (**Fig. 1e**). We then assessed whether *Chst9* is selective for the Grm8-MSN cluster using the Gini coefficient, a metric useful for identifying variables not equally shared across groups or clusters of cells. This analysis identified *Chst9* as a potential marker gene for the Grm8-MSN population, as it exhibits selectively high expression for that population only (**Fig. 1f**). The evaluation of *Chst9* expression within all other NAc cell populations, utilizing the same analysis pathway, confirmed this observation— every cluster outside of Grm8-MSNs lacked *Chst9* expression, identifying this as a unique marker gene (**Fig. 1g**).

### Validation of Chst9 as an Oprm1+ subpopulation marker gene

To validate snRNA-seq findings demonstrating that *Chst9* is in a distinct population of *Grm8*+/ *Oprm1*+ MSNs in the NAc, we employed RNAscope^39^, a widely implemented single molecule RNA fluorescence *in situ* hybridization (smRNA-FISH) platform for multiplexed transcript detection. Tissue sections containing the NAc from naïve rats were used for multiplexed hybridization using distinct probes for G*rm8*, O*prm1*, and C*hst9* (**Fig. 2a-c**). Consistent with rat NAc snRNA-seq data, G*rm8*, O*prm1*, and C*hst9* mRNA colocalized in a subset of NAc neurons (**Fig. 2b-c**).

**Figure 2.**
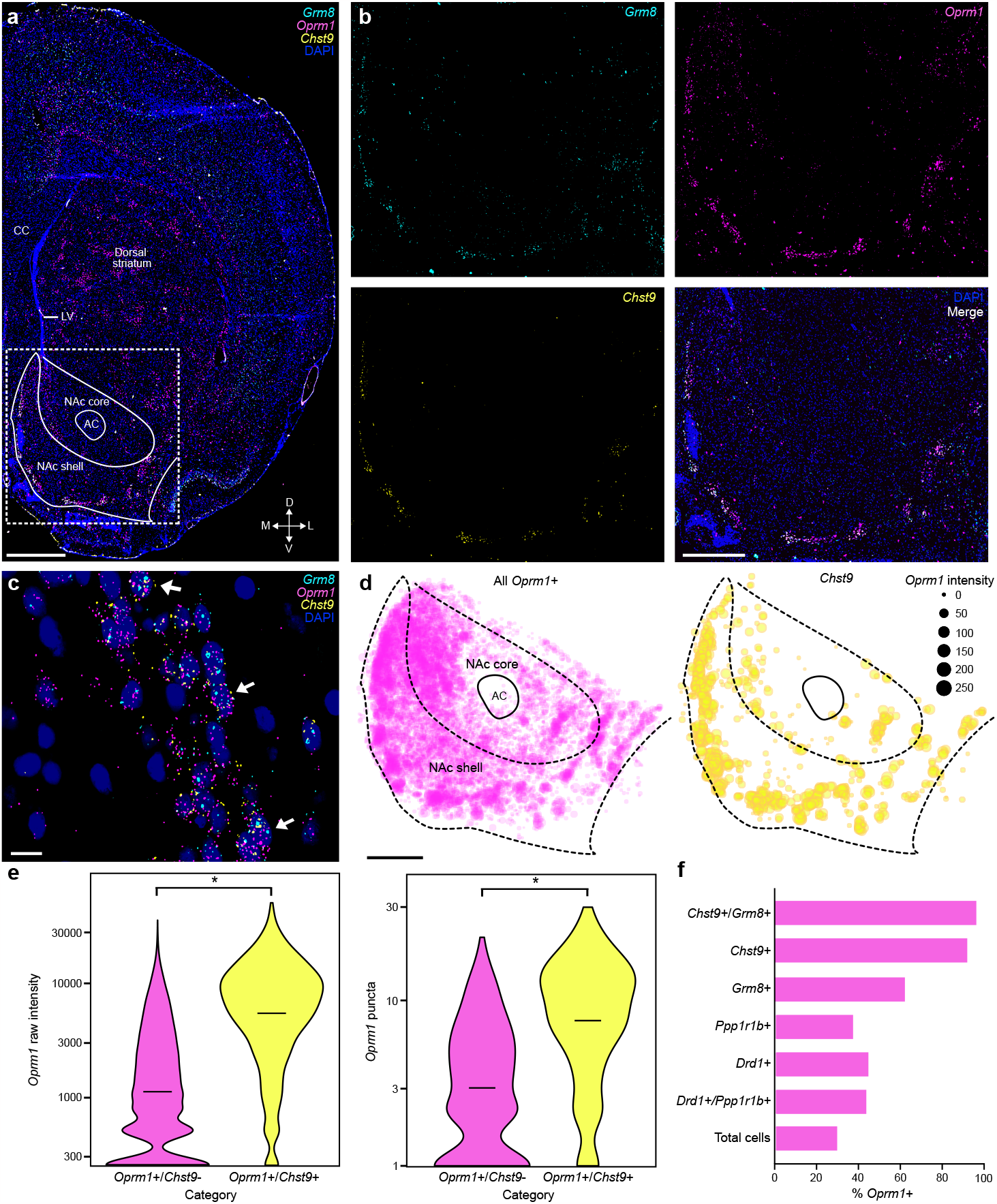
Validation of *Oprm1* subpopulation marker genes with multiplexed RNAscope single molecule RNA-FISH. **a**, Representative 4x image of *Grm8* (cyan), *Oprm1* (magenta), *Chst9* (yellow), and DAPI (blue) detection in a rat brain section with annotated anatomical landmarks. CC, corpus callosum; LV, lateral ventricle; AC, anterior commissure. Scale bar, 1 mm. **b**, Representative 20x images of *Oprm1, Grm8, Chst9*, and DAPI in the NAc. Scale bar, 500 μm. **c**, Identification of *Grm8*+/*Oprm1*+/*Chst9*+ cells (arrows) using probes targeting *Grm8, Oprm1*, and *Chst9*. Scale bar, 10 μm. **d**, Spatial distribution plots (n = 5 rats) of cells expressing *Oprm1* and *Chst9* mRNA. Scale bar, 500 μm. Mean *Oprm1* intensity per cell is represented in the spot area. **e**, Quantification and comparison of raw *Oprm1* intensity and puncta count in *Oprm1*+/*Chst9*+ and *Oprm1*+/*Chst9*-populations (nested t-test: p < 0.0001, p = 0.0002). **f**, Percentage of cells positive for *Oprm1* mRNA, classified within cells labeled by other MSN markers (or all NAc cells).

Additionally, we used smRNA-FISH to visualize the spatial distribution of targeted gene transcripts. Though *Oprm1* and *Grm8* expression can be observed across the broader NAc, *Chst9* expression was restricted to a much smaller group of cells, again highlighting the selectivity of this marker across NAc subpopulations (**Fig. 2c**). Curiously, this *Chst9*+ population appears to form discrete cellular clusters along the medial and ventral borders of the NAc shell subregion (**Fig. 2c**). To confirm this observation across all animals, region of interest (ROI) coordinates from quantified cell types were combined into an integrated spatial map, forming a spatially co-registered map of cell type location of the NAc (**Fig. 2d**). These maps further revealed that although Oprm1+ cells are distributed across the medial/lateral and dorsal/ventral axes of the NAc, *Chst9*+ cells were relatively confined along the NAc shell’s medial and lateral border (**Fig. 2d**). Additionally, by reflecting the degree of *Oprm1* expression in the area of each point, this map suggested that nearly all *Chst9*+ cells also abundantly expressed *Oprm1* (**Fig. 2d**).

To confirm the relative expression of *Oprm1* within different NAc cell types, we quantified the intensity and transcript-level puncta of *Oprm1* signal within all *Oprm1*+ cells. *Oprm1*+ cells were then further identified as *Chst9*+ or *Chst9*-. These analyses verified significantly elevated expression of *Oprm1* in *Chst9*+ cells as compared to *Chst9*-cells (**Fig. 2e**). We additionally evaluated the percentage of different NAc cell types, including *Drd1*+ and *Ppp1r1b*+ neurons, that express *Oprm1. Oprm1* was detected in in ∼40% of *Drd1*+ neurons, and in ∼30% of total NAc cells (**Fig. 2f**). However, *Oprm1* is expressed in over 90% of *Chst9*+ NAc neurons (**Fig. 2f**). In combination with snRNA-seq data, these observations confirm *Chst9* as a selective marker for NAc neurons that abundantly express *Oprm1* mRNA.

### Conservation of Chst9+ neurons across mammalian species

While these studies identify *Chst9*+ MSNs in the rat brain, we next investigated conservation of this NAc cell population across species, with a focus on higher-order mammals. To do this, we leveraged publicly available snRNA-seq atlases of the non-human primate (NHP)^36^ and human^38^ NAc that have previously been used to compare gene expression profiles and MSN subtypes across species^34^. As previously described, both NHP and human datasets identified unique MSN clusters that exhibit similar marker gene expression patterns to those observed in the rat NAc (**Fig. 3a-c**). The direct comparison of *Grm8, Oprm1*, and *Chst9* expression between these species revealed that one cell type in each respective dataset exhibited colocalization and abundant expression of both *Grm8/GRM8* and *Oprm1/OPRM1*. Notably, the same population was marked by cluster-selective *Chst9/CHST9* expression. While the rat literature had termed these populations Grm8-MSNs, NHP datasets termed them D1-NUDAP cells, and human datasets referred to these as MSN.D1_F cells (**Fig. 3a-c**).

**Figure 3.**
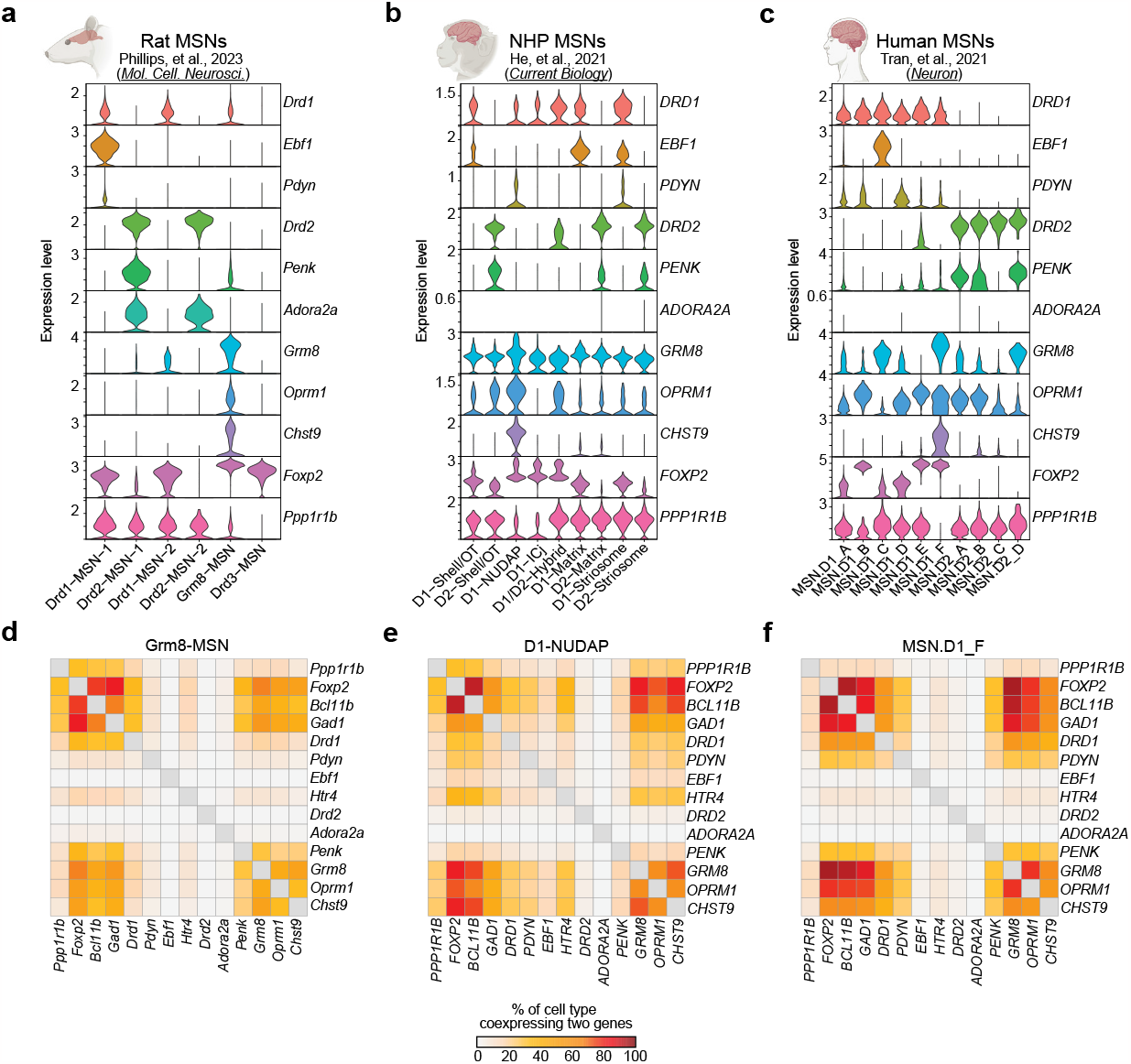
Identification of conserved populations of *Chst9+* MSNs in non-human primates and humans. **a-c**, Violin plots of MSN marker genes in transcriptionally defined cell populations of the rat (**a**), non-human primate (**b**), and human (**c**) NAc. **d-f**, Coexpression heatmaps showing the percent of cells within the rat Grm8-MSN cluster (**d**), non-human primate D1-NUDAP cluster (**e**), or human MSN.D1_F cluster (**f**) coexpressing identified genes.

Transcriptionally conserved cell types should be expected to display a similar enrichment and co-expression of other marker genes as well. To examine whether this was the case, we generated co-expression heatmaps of selected transcripts within Grm8-MSNs, D1-NUDAP cells, and MSN.D1_F cells (**Fig. 3d-f**). For this analysis, we prioritized previously characterized markers of MSNs or MSN subtypes, in addition to *Grm8, Chst9*, and *Oprm1*. Consistent with a potentially conserved role across rodent and primate NAc samples, we observed similar patterns of expression for additional marker genes in these populations. For example, each cluster included a high percentage of cells expressing the genes *Foxp2* and *Bcl11b*, commonly cited markers of MSNs^40,41^. Furthermore, an appreciable subset of cells in each cluster expressed *Drd1*/*DRD1* and *Pdyn*/*PDYN*, both established markers of D1-MSNs^42,43^. Additionally, feature expression plots from each species confirmed a similar expression of *Drd1/DRD1*, among other genes, across these three clusters (**Fig. S1**). Taken together, this finding demonstrates that Grm8-MSNs, and specifically *Chst9*+ neurons, are conserved across species and exhibit similar gene expression signaturesto previously described cell types in higher-order mammals.

## DISCUSSION

Recent large-scale single cell sequencing platforms have enabled comprehensive dissection of the molecular architecture of the striatum (including the NAc) in multiple species^32–36,38^. While these efforts have identified transcriptionally distinct subdivisions within larger D1-MSN and D2-MSN subclasses, the convergence of cell types and the reliability of specific marker genes across species remains unclear. Here, we leveraged published snRNA-seq datasets from the rat NAc to identify putative transcriptional markers of Grm8-MSNs, a poorly characterized population of neurons that has been identified in multiple rodent snRNA-seq datasets across different laboratories^33–35,44^. Notably, this cell population exhibits the highest expression of *Oprm1* within the NAc, and is selectively marked by mRNA for the carbohydrate sulfotransferase gene *Chst9*. At an anatomical level, *Chst9*+ MSNs populate the ventral and medial borders of the NAc shell subregion, where they localize in discrete clusters that co-express *Grm8*. Moreover, we demonstrate that *Chst9*+ cells are observed in NAc subpopulations in the monkey and human NAc, providing evidence for a conserved transcriptional profile. Together, these results help to define a poorly characterized cell type in the NAc, and highlight the need for further investigation into the circuit and behavioral role of *Chst9*+ MSNs.

The anatomical localization of *Chst9*+ MSNs at the border of the rat NAc may generate confusion given the proximity to other nearby landmarks, including the Islands of Calleja and anterior components of the ventral pallidum that perforate the ventral NAc shell. Fortunately, these distinct nuclei can be reliably distinguished from principal NAc neurons at the molecular level. For example, the Islands of Calleja are densely organized pockets of granule cells within the ventral striatum, which play a key role in grooming behavior^45^. While the Islands of Calleja can easily be localized in a coronal section based on cell density alone (e.g., increased DAPI+ territories in **Fig. 2a**), they are also marked by robust expression of the *Drd3* receptor^45^, which is largely absent in *Chst9*+ MSNs. Likewise, *Chst9*+ MSNs do not express other markers of ventral pallidum neuronal subtypes, including *Npas1* and *Pvalb*^46,47^, but do express appreciable levels of D1-MSN markers including *Drd1, Bcl11b, Pdyn*, and *Ppp1r1b*. Together with molecular conservation in higher-order mammals, these results argue that *Chst9*+ MSNs represent a distinct cell type that is both transcriptionally and anatomically separate from other defined MSN subtypes^34,36^.

Strong evidence supports a critical role of μORs in the NAc in the rewarding and reinforcing properties of opioids. However, the specific contributions of distinct cell types or anatomical subregions is only beginning to be understood. Landmark studies previously identified “hotspots” for opioid effects on hedonic reactions to sucrose reward, as defined by enhancement of orofacial affective taste reactivity measures^8,11,48,49^. These studies outlined a rostrodorsal domain of the medial NAc shell as having the most robust effects in stimulating “liking” of palatable rewards, whereas μOR activation in other regions of the NAc produced more moderate effects or even generated food avoidance. Notably, the strongest hedonic effects of μOR activation at least partially overlap with the distribution of *Chst9*+ clusters in the NAc shell (**Fig. 2d**), suggesting a possible role for these cells in hedonic reactions to palatable foods. Given the abundant expression of *Oprm1* in *Chst9*+ cells and the high fraction of *Chst9*+ cells that express *Oprm1*, it would be a reasonable assumption that these cells would be among the most responsive to μOR activation. Prior evidence from *Fos* plume mapping following direct NAc microinfusions of μOR agonists identified dose-dependent elevations in FOS protein^11,14,49^, but the molecular identity of FOS+ cells was not investigated. Similarly, an early single cell RNA-seq investigation revealed increased activity-driven gene expression in a small subset of D1-MSNs after morphine exposure in a mouse model^50^, although it is unclear if these changes were confined to a distinct subset of D1-MSNs. Likewise, a recent study found no change in *Fos* (or related AP-1 transcription factor family member *Fosb*) in Grm8-MSNs following either single or repeated cocaine administration^34^. Future studies will be needed to comprehensively define the electrophysiological and transcriptional responses of *Chst9*+ cells to opioids and other psychoactive drugs.

Although we identified *Chst9* as the most specific marker for Grm8-MSNs, we also detected elevated levels of other potential marker genes in this population (**Fig. 1e**). These included *Tshz1* (Teashirt zinc finger homeobox 1) and *Tac3* (Tachykinin precursor 3; the rat ortholog of mouse *Tac2*). Each of these genes have recently been discovered by other groups to mark specific populations of MSNs in the dorsal and ventral striatum, and cells expressing these genes have been shown to be important for aversive aspects of motivated behavior^43,51^. *Tshz1* is expressed by a subgroup of striosomal D1-MSNs located in the dorsal striatum, but is also found in the ventral striatum. Notably, *Tshz1*+ neurons in the dorsal striatum are activated by aversive stimuli, and optogenetic stimulation of these cells causes place aversive and suppresses locomotion^43^. Similarly, inhibiting *Tshz1*+ MSNs impairs learning related to punishment, but does not impact reward learning. However, while this study characterized *Tshz1*+ MSNs in the dorsal striatum, nothing is known about the behavioral role of *Tshz1*+ neurons in the ventral striatum or NAc, and we did not observe significant *Chst9* expression in the dorsal striatum. Similarly, another recent study identified *Tac2* mRNA to mark a subpopulation of D1-MSNs in the mouse NAc^51^. This study found that acute cocaine decreased activity of *Tac2*+ neurons, and stimulation of this population suppressed cocaine place preference and self-administration. Given the enrichment of *Tac3* in rat Grm8-MSNs, the cells targeted in this mouse study likely overlap to some extent with the *Chst9*+ cells investigated in the present manuscript. However, further work is needed to characterize the circuit and behavioral role of this population, including synaptic inputs and downstream targets.

In summary, we have identified a transcriptionally and anatomically unique population of NAc MSNs marked by high expression of *Oprm1* and selective expression of *Chst9*. Given that this population expresses other markers of D1-MSNs, it is likely that prior labeling studies or genetic manipulations may have targeted this cell population along with more classically defined D1-MSNs that do not express *Chst9* and exhibit lower expression of *Oprm1*. For example, use of *Drd1* or prodynorphin (*Pdyn*) gene promoter sequences is a common strategy for cell-restricted expression of transgenes in D1-MSNs^2,21,52–54^. Using this strategy, prior work has revealed that *Oprm1* expression in *Pdyn*+ cells (using a *Pdyn* BAC transgenic mouse) is sufficient to restore opioid reward in an *Oprm1*^-/-^ mouse^10^. However, based on the present results (as well as recent work demonstrating that *Pdyn* regulatory elements are accessible in Grm8-MSNs^55^), it is likely that this manipulation restores *Oprm1* expression in multiple MSN subtypes including *Chst9*+ cells, which may each contribute to opioid reward in distinct ways. Our results highlight the previously underappreciated diversity within broad striatal MSN subclasses, and underline the need for continued exploration of the distinct roles of unique cell types.

## METHODS

### Animals

Adult male and female Sprague-Dawley rats were obtained from Charles River Laboratories (Wilmington, MA, USA). Rats were cohoused in pairs in plastic filtered cages with nesting enrichment in an Association for Assessment and Accreditation of Laboratory Animal Care–approved animal care facility maintained between 22° and 24°C on a 12-hour light/12-hour dark cycle with ad libitum food (Lab Diet Irradiated rat chow) and water. Bedding and enrichment were changed weekly by animal resources program staff. A total of 6 naïve rats (n = 3M/3F) were used in initial *Chst9* population and MSN characterization experiments. Animals were randomly assigned to experimental groups. All experiments were approved by the University of Alabama at Birmingham Institutional Animal Care and Use Committee (IACUC).

### Brain tissue collection

Rats were euthanized by live decapitation using a guillotine (World Precision Instruments, Sarasota, FL, USA). Brains were rapidly removed and submerged in 2-methylbutane, chilled on dry ice, for approximately 30 seconds. Flash-frozen brains were then wrapped in aluminum foil and placed on dry ice. Tissue was stored at -80°C until the day of sectioning. After a 30-minute equilibration at -20°C, 10 μm coronal sections were obtained using a Leica CM 1850 cryostat (Deer Park, IL, USA) set to -20°C. Sections containing the NAc, ranging from 1.68-1.92 mm anterior-posterior (A/P) to Bregma, were mounted onto room temperature frosted microscope slides, air dried for 60 minutes at -20°C, and stored at -80°C until staining.

### Tissue preparation and imaging for RNAscope

The RNAscope Multiplex Fluorescent v2 assay kit (ACD Bio, 323110) and RNAscope 4-plex ancillary kit (ACD Bio, 323120) were used to stain sections, following the manufacturer’s recommended protocol. Probe sets used include DAPI (320858), Rn-Oprm1-C1 mRNA (410691-C1), Rn-Grm8-C2 mRNA (1149671-C2), Rn-Chst9-C4 mRNA (1149681-C4), Rn-Ppp1r1b-C2 mRNA (1048941-C2), and Rn-Drd1-C3 mRNA (317031-C3). Three different groups of sections were probed, with varying marker probe combinations— group 1 (*Chst9* population validation experiment) probes included *Oprm1, Grm8*, and *Chst9*; group 2 (MSN marker experiment) probes included *Oprm1, Ppp1r1b*, and *Drd1*; group 3 (MSN marker experiment) probes included *Oprm1, Ppp1r1b*, and *Chst9*. Channels were matched with genes according to relative expression level and channel background in different experiments: the green channel (520 nm dye, recommended for high expressors) was assigned to *Grm8, Ppp1r1b*, and *Oprm1*, the red channel (570 nm dye, recommended for low expressors) was assigned to *Oprm1*, and the white channel (690 nm dye, recommended for low expressors) was assigned to *Chst9* and *Drd1*. 5 total sections (1 each from 3M/2F) were used for the *Chst9* population validation experiment, and 6 total sections (1 each from 3M/3F) were used for each of the MSN marker experiment. After staining, sections were imaged using the BZ-X800 Keyence Microscope, and then stitched using the BZ-X800 Keyence Analyzer software. High sensitivity images were taken using OP-87767 filter cubes for GFP (Channel 1, green), Texas Red (PE) (Channel 2, red), DAPI (Channel 3, blue) and Cy5 (Channel 4, white) at 4x (PlanApo NA 0.20), 20x (Plan Fluor NA 0.45 Ph1), and 100x (PlanApo 1.45/0.113 mm Oil) magnifications.

### Image analysis

Approximate A/P coordinates of sections were determined by comparing anatomical landmarks within 4x whole section images to those in coordinate and region-labeled diagrams of the Paxinos and Watson rat brain atlas^56^. These A/P distinctions were used to process 20x ROI images of the NAc. This protocol was used for the *Chst9* population validation and MSN marker experiments. To isolate the NAc for quantification, brain atlas templates at the corresponding A/P coordinate to that of a given section were overlaid on 20x images of that section in Adobe Illustrator (San Jose, CA, USA) using anatomical landmarks as a guide. This overlay image was then imported into FIJI^57^ to create NAc-specific ROIs for that 20x image. These ROIs were then applied to each individual channel of the original 20x image, and the NAc was cropped from the rest of each channel’s image using the Clear Outside plugin. DAPI channel nuclei-specific ROIs were generated with the StarDist-2D plugin^58^, using the “Versatile (fluorescent nuclei)” model with a probability/score threshold of 0.20 and an overlap threshold of 0.45. Nuclei-specific ROIs were then applied to images containing *Oprm1, Grm8, Chst9, Ppp1r1b*, or *Drd1* probe signal, and raw signal intensity within each ROI was measured. Cell types were categorized based on positive marker colocalization within a single ROI, and intensity values were determined within different cell types, using R Studio. Nuclei-specific ROI positions were used to plot locations of different cell types, using R Studio.

100x magnification images were taken using the z-stack setting. One image for each channel within a stack was chosen for analysis, for a total of 10-14 images per animal within the *Chst9* population validation experiment. Selected channel images for one total z-stack image were merged, and the resulting overlay was converted to Nikon (.nd2) format using NIS-Elements software. Nuclei-specific ROIs were generated around DAPI staining using QuPath^59^ software version 0.3.2. Cell detection parameters were set to background radius 0 px, median filter radius 0 px, minimum area 100 px^2, maximum area 3000 px^2, threshold 35, and cell expansion 3 px. For cell detection, “split by shape”, “include cell nucleus”, “smooth boundaries”, and “make measurements” boxes were checked. Marker puncta-specific ROIs were then generated, and counts were measured within each DAPI ROI for an image. Subcellular detection parameters were set to expected spot size 12 px^2, minimum spot size 0.5 px^2, and maximum spot size 25 px^2. Cell types were categorized based on positive marker colocalization within a single ROI, and puncta counts were determined within different cell types, using R Studio.

### Statistical analysis

All image data were analyzed statistically with Prism software using the nested t-test (*Chst9* population validation experiment) or Welch’s t-test (fentanyl experiment). A 95% confidence level was set for every test, with a definition of statistical significance at p < 0.05. All graphs, diagrams, and image edits were made using Prism, Adobe Illustrator, R Studio, and FIJI.

### Published snRNA-seq data analysis

Data objects for NHP and human NAc snRNA-seq datasets were obtained from data repositories outlined in each respective publication. The data object for the rat NAc snRNA-seq dataset was provided directly by Robert Phillips at the University of Alabama at Birmingham (UAB). All data objects were analyzed with R software. Data for *Grm8, Oprm1, Chst9, Ppp1r1b*, and *Drd1* expression were obtained in the form of UMAP cluster plots and violin plots through Seurat^60^, ggplot2, and dplyr package processing in R Studio.

## DATA AVAILABILITY

All relevant data that support the findings of this study are available by request from the corresponding author (J.J.D.).

## ACKNOWLEDGEMENTS

We thank all current and former Day Lab members for assistance and support. This work was supported by NIH grants DP1DA039650, R01MH114990, R01DA053743, R01DA054714, and the McKnight Foundation Neurobiology of Brain Disorders Award (JJD). R.A.P.III is supported by the AMC21 scholar program and the UAB T32 in the Neurobiology of Cognition and Cognitive Disorders (T32NS061788).

## AUTHOR CONTRIBUTIONS

E.A. and J.J.D. conceived of experiments. E.A. performed all experiments. E.A. and J.J.D. harvested brains for the marker validation experiment. E.A. analyzed smRNA-FISH data with assistance from R.A.P.III and K.L.B. R.A.P.III analyzed published snRNA-seq data. All authors have approved the final version of the manuscript.

---

**Figure S1.**
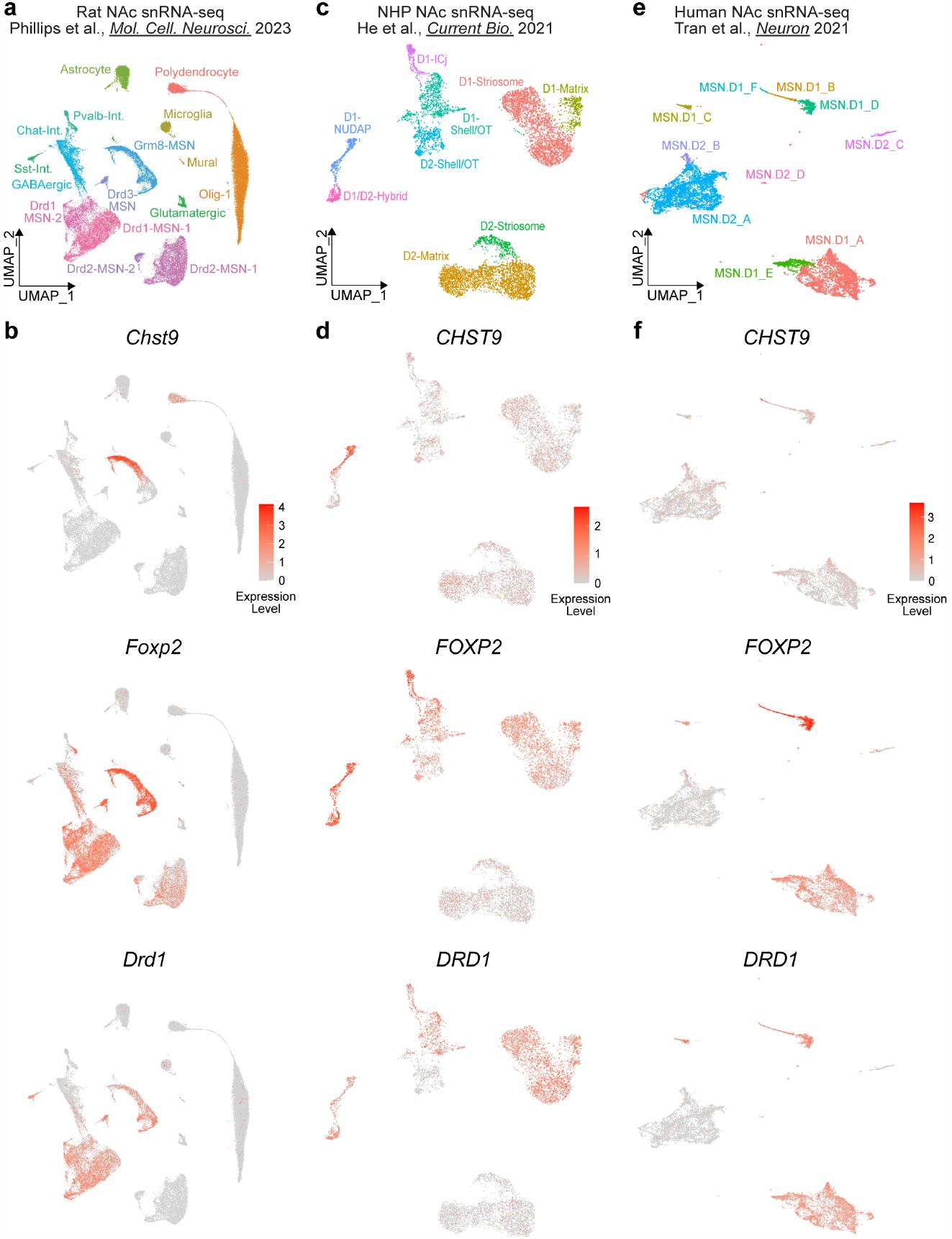
Comparison of cell type marker genes across species, **a**, UMAP of previously published snRNA-seq data from NAc of male and female Sprague-Dawley rats, **b**, Rat NAc UMAP plots, from top to bottom: *Chst9* cluster plot, *Foxp2* cluster plot, and *Drd1* cluster plot, c, UMAP of previously published snRNA-seq data from the striatum of non-human primates, **d**, NHP striatum UMAP plots, from top to bottom: *CHST9* cluster plot, *FOXP2* cluster plot, and *DRD1* cluster plot, **e**, UMAP of previously published snRNA-seq data from the striatum of humans, **f**, Human striatum UMAP plots, from top to bottom: *CHST9* cluster plot, *FOXP2* cluster plot, and *DRD1* cluster plot.

